# *In silico* and *In vitro* Investigations of Antihelmintic Activities of some Selected Approved Drugs

**DOI:** 10.1101/837427

**Authors:** Christopher Arinze Agube, Daniel Lotanna Ajaghaku, Ikemefuna Chijioke Uzochukwu

## Abstract

The search for alternative methods to mitigate the drawbacks associated with wet laboratory drug discovery has been a major challenge in drug design and has limited the options available in the fight against most Neglected Tropical Diseases (NTDs) such as helminthiasis. We investigated the binding affinities of some approved drugs to *Ascaris suum* mitochondrial rhodoquinol fumarate reductase enzyme (MRFR),an essential enzyme for *ascaris* survival, and the possibility of repurposing these drugs as antihelmintic agents using *in silico* molecular docking and *in vitro* paralysis and mortality times of fifteen selected front runners.Two hundred approved drugs were selected from ZINC^®^ database based on bioactivity scores while MRFR(PDB code, 3vra) was obtained from the Protein Data Bank (PDB). Both were prepared using AutoDock tools v.1.5.6 and Chimerav.1.9.The docking protocol was validated by computationally reproducing the binding of atpenin to MRFR. The selected approved drugs and the receptor were docked using AutoDockVina v. 4.0. The docking results were analyzed using PyMoL v. 1.4.1.The paralysis and mortality times of the identified frontrunners against *Pheretima posthuma* were determined *in vitro* and synergistic testings were done by the checkerboard method. Fifteen drugs had binding free energies between −7.825 to 11.025 kcal/mol while four of these drugs (mefloquine, doxycycline, mepacrine and proguanil) emerged as major frontrunners by both *in silico* and *in vitro* assessments. The paralysis and mortality times of the four drugs were between 0.33-0.50 hr as against 1.80-2.36 hr for albendazole. They were therefore predicted to have ability to affect MRFR in the same manner as atpenin hence, suggestive of potential antihelmintic activity. *In vivo* investigation of these frontrunner drugs is strongly recommended.

**Author Summary:** For decades, intestinal worm infections have been a major public health problem particularly in the tropical regions of the world with infection figures put at 5.3b [16].

This disease is often associated with poor hygiene and sanitation hence is predominant in the resource-poor nations with physical infrastructural deficit [8].

Children between the ages of 2-10 years are the most vulnerable by reason of their hand habits with morbidity leading to compromise of physical and cognitive development while several sufferers have had to live disability-adjusted life years as a consequence [12].

Efforts at implementing improved hygiene and mass deworming campaigns have not delivered the intended outcomes with many reports coming from clinicians regarding gradual development of resistance to the standard treatments [9].

Given the huge cost and time invested in traditional drug discovery and the rarity of this disease in regions of the world with the resources for research, no new drugs have been developed in the past two decades [18].

The advent of computational techniques in drug discovery has come to mitigate these drawbacks. This study exploited this technique to full effect and through laboratory assays and clinical investigation of selected approved drugs, mepacrine and doxycycline were identified as potential antihelmintic agents fit for combination therapy.

## Introduction

The fortuitous nature, enormous cost and huge time invested in traditional drug discovery and development is a major factor that has fueled the inertia in most pharmaceutical companies to engage in research into new chemical entities. This is a consequence of the unpredictability of these researches hence results often do not justify the effort.

This scenario has mostly affected the neglected tropical and orphan disease domains. It is so due to the limited number of sufferers and the demography that stratifies them to the resource-poor nations [13].

As a fall out of this, pharmaceutical majors located in regions with the technology and wherewithal do not find it attractive to direct research into these areas due to marginal profit prospects.

Helminthiasis, a major Neglected Tropical Disease (NTD) has long been identified by the World Health Organization (WHO) as a disease with very high morbidity rate and cognitive deficit especially in school-aged children [20, 3]. The prevalence is mostly confined to the tropical and subtropical regions where there still exists infrastructural deficit; poor sanitation; use of untreated fecal matter as fertilizers and subsisting bare soil defecation [1].

In addition, no new antihelmintic agents have been added to the list in the past decade with gradual development of resistance to the standard treatments being reported [17, 15].

Drug repurposing has recorded major milestones in several treatment landscapes hence this technique has been considered a veritable tool to achieve this objective in antihelmintic sphere.

It is a well-known fact that observed pharmacological activity of drugs is premised on their ability to form stable complexes with their receptors while the magnitude of activity is a function of avidity of binding. The application of computational techniques in addition to *in vitro* tools in modern day drug design has afforded the benefits of specificity and druglikeness optimization hence mitigation of cost and time investment.

Molecular docking simulations represent the classical method in computational studies and the successes recorded using this technique both in antihelmintic and other drug discovery landscapes gave impetus for the employment of this technique in this study. Prominent among such studies is the work of Uzochukwu, *et al.*, [18] which revealed the potentials of some approved drugs as antihelmintic agents.

This study investigated the antihelmintic potential of some approved drugs using *Ascaris suum* mitochondrial rhodoquinol fumarate reductase enzyme as target. The *in vitro* paralysis and mortality times of the frontrunners using the *ascaris* surrogate, *Pheretima posthuma* were determined.

## Results

On the basis of bioactivity scores similarity of the approved drugs to atpenin, the in-house drug database was sorted and one hundred drugs were selected. And the docking protocol was validated by superimposition of the experimental atpenin A5 on the atpenin from docking which showed a near perfect fit (Fig 1).

**Fig. 1:**
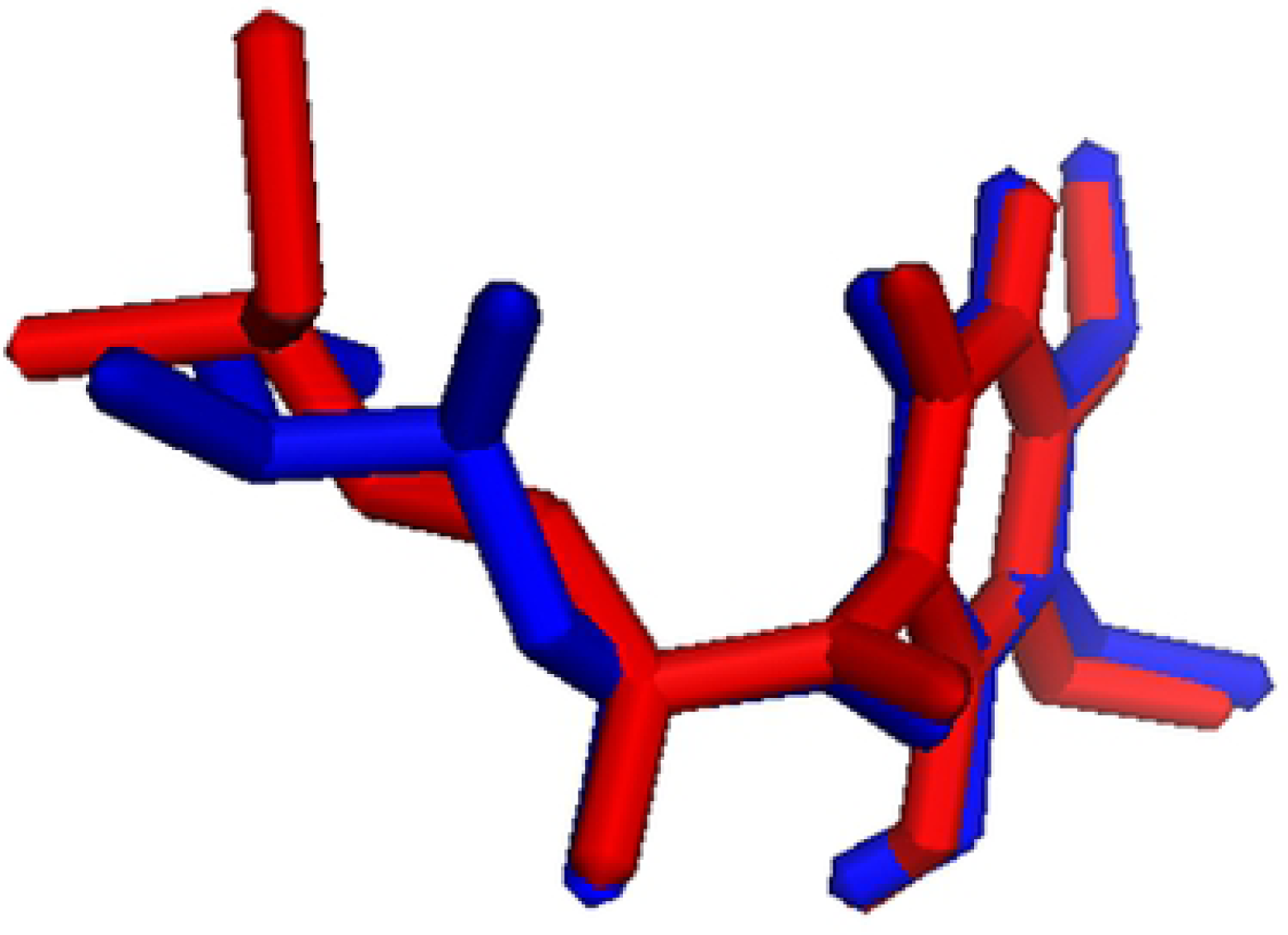
The superimposition of the experimental (red) and docked atpenin (blue)

The post docking analysis yielded two hundred drugs(including their isomers) with binding energies within the range of atpenin (–7.825 kcal/mol).

Out of this list, fifteen drugs with the closest binding energies to atpenin were selected as frontrunners.

This result is presented in Table 1.

**Table 1:** Binding energy and molecular descriptors of frontrunner drugs.

| Drug | Binding Energy (kcal / mol± SEM) | MW (g / mol) | XlogP | tPSA($\text{\AA}$) | Solubility in water | Protein Binding(%) |
| --- | --- | --- | --- | --- | --- | --- |
| Glimepiride | $-11.400 \pm 1.200$ | 490.62 | 1.8 | 130 | Freely soluble | 99.5 |
| Mefloquine | $-11.023 \pm 0.050$ | 378.31 | 3.67 | 45.15 | Poorly soluble | 98 |
| Mebendazole | $-9.825 \pm 0.640$ | 295.29 | 2.95 | 84.1 | Very slightly soluble | 90.95 |
| Carvedilol | $-9.775 \pm 0.320$ | 406.47 | 4.04 | 75.75 | Soluble | 98 |
| Doxycycline | $-9.600 \pm 0.000$ | 444.43 | 0.196 | 181.62 | Soluble | 95 |
| Oxazepam | $-9.50 \pm 0.000$ | 286.71 | 2.31 | 61.69 | Soluble | 80 – 99 |
| Praziquantel | $-9.025 \pm 1.120$ | 312.40 | 3.36 | 40.62 | Soluble | 80 – 85 |
| Mepacrine | $-8.975 \pm 0.850$ | 399.95 | 6.72 | 37.39 | Sparingly soluble | 80 -90 |
| Nalidixic Acid | $-8.425 \pm 0.490$ | 232.23 | 1.02 | 70.50 | Soluble | 93 |
| Metronidazole | $-8.350 \pm 0.340$ | 171.15 | -0.46 | 81.19 | Soluble | 20 |
| Chloroquine | -8.325 ± 0.430 | 319.87 | 5.06 | 28.16 | Soluble | 55 |
| Albendazole | -8.200±0.080 | 265.33 | 1.70 | 84.08 | Poorly soluble | 70 |
| Proguanil | -8.000± 0.000 | 253.73 | 2.53 | 83.79 | Soluble | 75 |
| Atpenin (Probe) | -7.825 ± 0.520 | 366.24 | 2.64 | 84.86 | Insoluble | NA |
| Pyrimethamine | -7.775 ± 0.580 | 248.71 | 3.0 | 77.82 | Soluble | 87 |
| Tinidazole | -7.400± 0. 670 | 247.27 | -0.32 | 97.78 | Soluble | 12 |

Four out of the fifteen frontrunner drugs were found to exploit the same binding pocket and interacted with the same amino acid residues within the receptor site (Figs 2 and 3).

**Fig 2:**
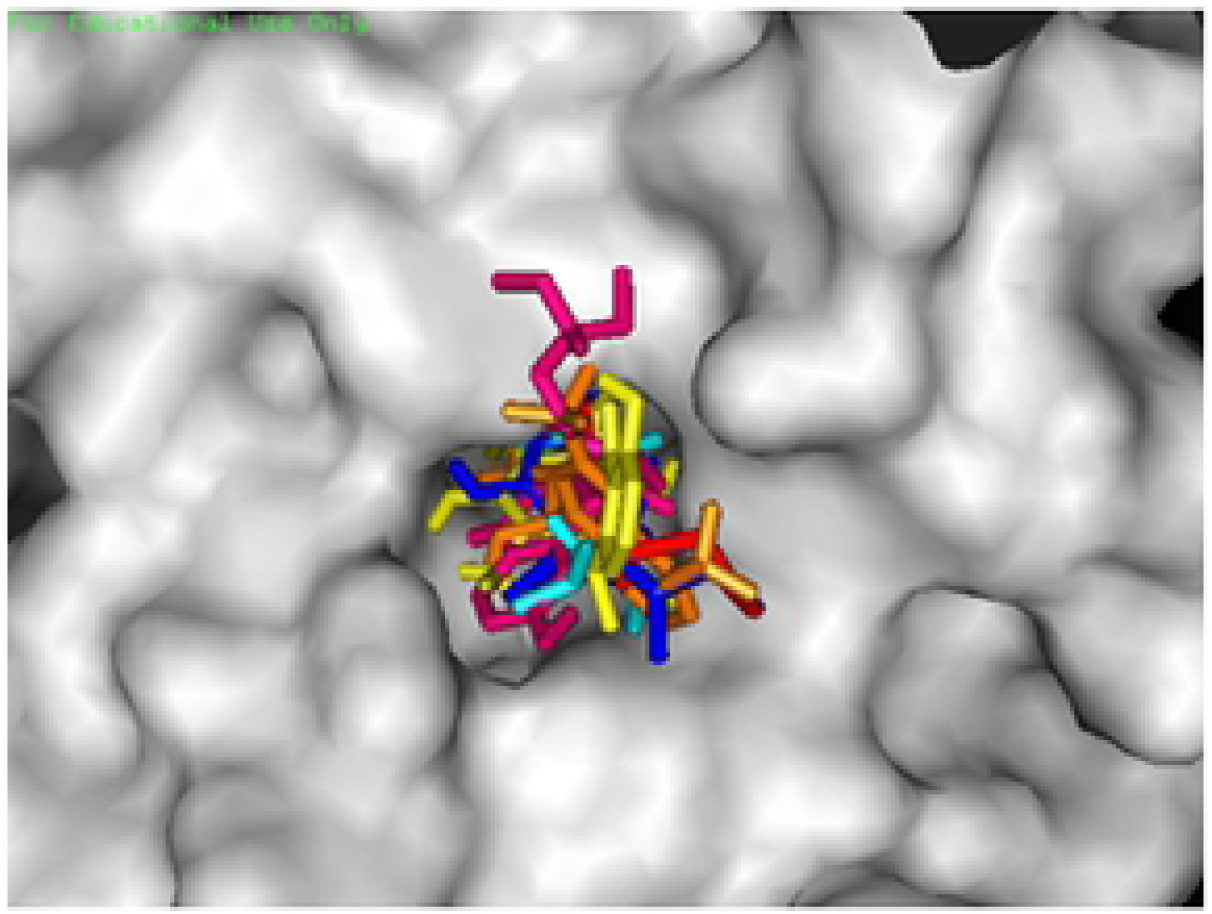
Docking pose of the binding pocket exploited by the four frontrunners and the probe (atpenin)

**Fig. 3:**
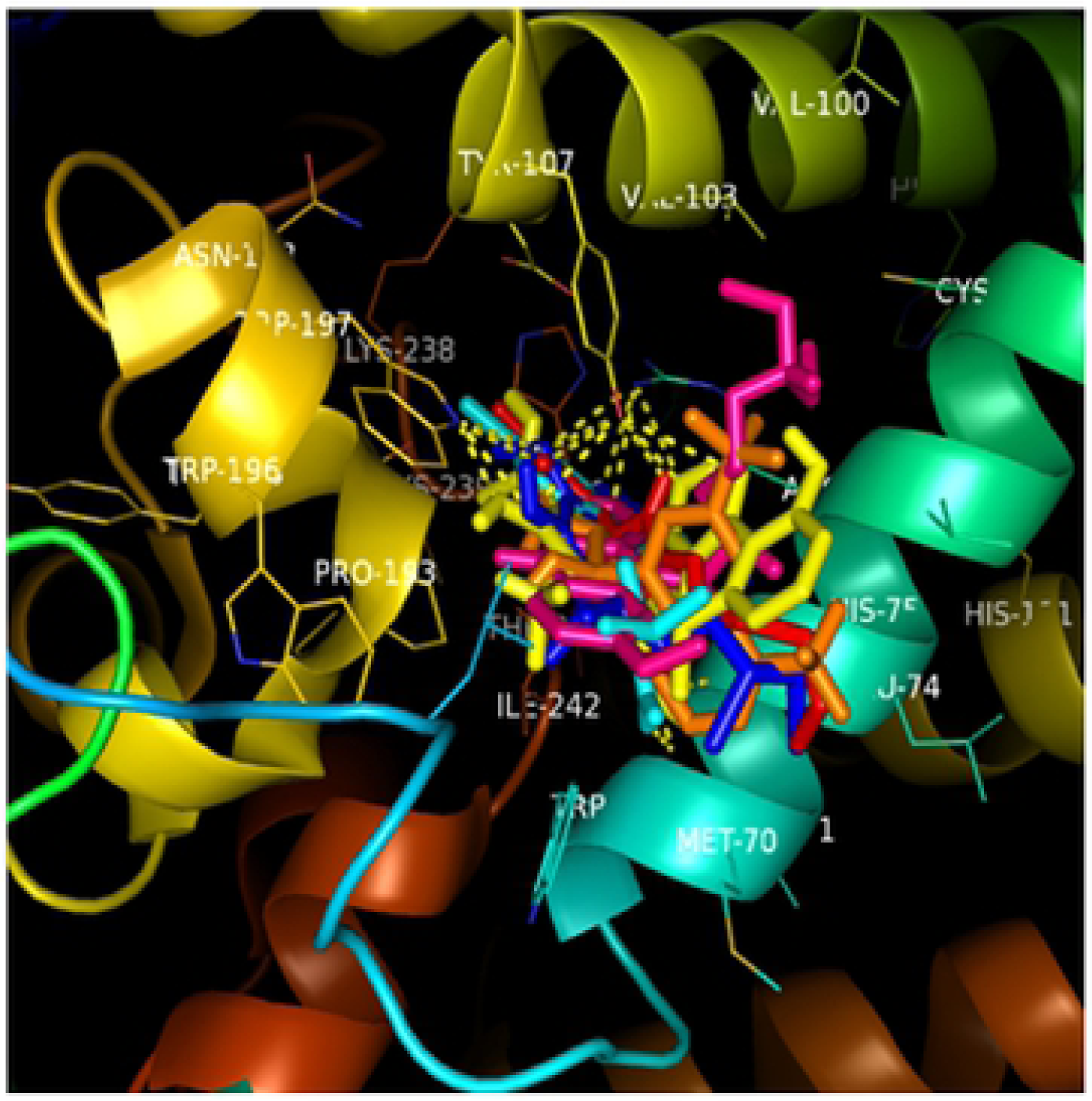
The Interactions between frontrunners and amino acids residues of the target.

The *in vitro* antihelmintic evaluation of the frontrunners across the seven assay points was dose-dependent.

Four drugs showed the best activity in this respective decreasing order: mefloquine, doxycycline, proguanil and mepacrine. The asterisk sign represents the gradation of activity hence triple asterisk represents the highest activity while the double and single asterisk represent the next in ranking order. This paralysis and mortality time is presented in Tables 2 and 3.

**Table 2:** Mean Paralysis Time (hr) ± SEM of *Pheretima posthuma*.

| Drugs | Concentration (mg/ml) |  |  |  |  |  |  | Mean Weight of Worm |
| --- | --- | --- | --- | --- | --- | --- | --- | --- |
|  | 5.0 | 2.50 | 1.25 | 0.625 | 0.3125 | 0.15625 | 0.07825 |  |
| Nalidixic acid | 3.30 ± 0.18 | 4.24 ± 0.10 | 4.9 ± 0.11 | 5.21 ± 0.22 | 7.77 ± 0.30 | 8.47 ± 0.24 | 9.16 ± 0.38 | 0.74 ± 0.14 |
| Metronidazole | 2.71 ± 0.34 | 3.44 ± 0.12 | 4.32 ± 0.42 | 4.78 ± 0.21 | 5.42 ± 0.15 | 6.71 ± 2.08 | 8.08 ± 1.25 | 0.75 ± 0.08 |
| Pyrimethamine | 2.33 ± 0.40 | 2.56 ± 0.00 | 2.63 ± 0.08 | 2.85 ± 0.01 | 4.31 ± 0.08 | 4.76 ± 0.10 | 5.29 ± 0.21 | 0.76 ± 0.09 |
| Chloroquine | 1.43 ± 0.98 | 1.47 ± 1.06 | 1.50 ± 0.60 | 1.51 ± 0.68 | 2.57 ± 0.17 | 3.41 ± 1.14 | 4.32 ± 0.39 | 0.73 ± 0.10 |
| Carvedilol | 1.29 ± 0.68 | 1.40 ± 0.61 | 1.57 ± 0.31 | 1.97 ± 0.00 | 2.29 ± 0.15 | 2.78 ± 0.95 | 2.84 ± 0.95 | 0.72 ± 0.03 |
| Praziquantel | 1.96 ± 0.00 | 1.99 ± 0.59 | 2.62 ± 0.18 | 2.81 ± 0.08 | 3.41 ± 0.13 | 3.76 ± 1.45 | 4.30 ± 0.92 | 0.78 ± 0.11 |
| Albendazole | 1.91 ± 0.00 | 1.91 ± 0.00 | 3.85 ± 0.08 | 4.18 ± 0.12 | 4.88 ± 0.28 | 6.39 ± 0.96 | 6.95 ± 1.52 | 0.73 ± 0.13 |
| Tindazole | 1.83 ± 0.00 | 1.83 ± 0.00 | 3.21 ± 0.01 | 3.71 ± 0.37 | 5.00 ± 0.51 | 5.40 ± 0.17 | 6.12 ± 0.21 | 0.79 ± 0.08 |
| Mebendazole | 1.80 ± 0.00 | 1.92 ± 0.21 | 1.92 ± 0.09 | 2.05 ± 0.16 | 3.79 ± 0.12 | 4.64 ± 1.06 | 4.88 ± 1.02 | 0.75 ± 0.09 |
| Glimepiride | 1.32 ± 0.17 | 1.35 ± 2.31 | 1.67 ± 0.08 | 1.78 ± 0.05 | 2.88 ± 0.18 | 3.24 ± 1.09 | 4.25 ± 1.29 | 0.73 ± 0.05 |
| Oxazepam | 0.84 ± 0.34 | 1.09 ± 0.86 | 1.21 ± 0.55 | 2.05 ± 0.03 | 2.25 ± 0.00 | 2.45 ± 1.85 | 2.95 ± 1.06 | 0.75 ± 0.06 |
| Mepacrine* | 0.50 ± 0.03 | 0.52 ± 0.00 | 0.70 ± 0.19 | 0.84 ± 0.24 | 1.35 ± 0.69 | 1.67 ± 0.88 | 1.93 ± 0.85 | 0.77 ± 0.06 |
| Proguanil* | 0.45 ± 0.03 | 0.53 ± 0.08 | 0.61 ± 0.34 | 0.81 ± 0.33 | 1.81 ± 0.34 | 2.07 ± 1.05 | 2.16 ± 1.03 | 0.74 ± 0.04 |
| Doxycycline ** | 0.39 ± 0.00 | 1.09 ± 0.08 | 1.33 ± 0.70 | 1.71 ± 0.20 | 1.91 ± 0.11 | 2.01 ± 0.93 | 2.02 ± 1.02 | 0.76 ± 0.08 |
| Mefloquine*** | 0.33 ± 0.00 | 0.34 ± 0.00 | 0.39 ± 0.08 | 0.50 ± 0.04 | 0.56 ± 0.02 | 1.05 ± 0.60 | 1.57 ± 0.77 | 0.72 ± 0.05 |
| Control > 12.00 | | | | | | | | 0.76 ± 0.05 |

**Table 3:** Mean Mortality Time (hr) ± SEM of *Pheretima posthuma*.

| Drugs | Concentration (mg/ml) |  |  |  |  |  |  | Mean weight of worms (g) |
| --- | --- | --- | --- | --- | --- | --- | --- | --- |
|  | 5.00 | 2.50 | 1.25 | 0.625 | 0.3125 | 0.15625 | 0.07825 |  |
| Nalidixic acid | 3.89 ± 0.40 | 5.74 ± 0.44 | 6.18 ± 0.32 | 6.94 ± 0.47 | 8.97 ± 0.96 | 10.26±0.70 | 11.18±1.22 | 0.74 ± 0.14 |
| Pyrimethamine | 2.87 ± 0.22 | 3.03 ± 0.53 | 3.89 ± 0.18 | 4.17 ± 0.85 | 5.29 ± 0.73 | 6.11 ± 0.72 | 6.85 ± 0.36 | 0.76 ± 0.09 |
| Metronidazole | 2.89 ± 0.13 | 3.73 ± 0.35 | 4.86 ± 0.43 | 5.55 ± 0.70 | 6.94 ± 0.9 | 7.82 ± 0.95 | 9.87 ± 1.32 | 0.75 ± 0.08 |
| Praziquantel | 2.43 ± 0.29 | 2.67 ± 0.69 | 3.51 ± 0.21 | 3.98 ± 0.55 | 4.74 ± 0.32 | 6.12 ± 0.86 | 7.86 ± 1.16 | 0.78 ± 0.01 |
| Albendazole | 2.36 ± 0.36 | 2.59 ± 0.49 | 4.03 ± 0.55 | 5.17 ± 0.46 | 6.23 ± 0.40 | 7.85 ± 1.04 | 8.19 ± 0.85 | 0.73 ± 0.13 |
| Mebendazole | 2.19 ± 0.52 | 2.40 ± 0.82 | 2.56 ± 0.72 | 2.92 ± 0.41 | 4.68 ± 0.81 | 6.21 ± 0.66 | 6.88 ± 1.48 | 0.75 ± 0.09 |
| Tindazole | 2.30 ± 0.26 | 2.46 ± 0.31 | 3.84 ± 0.58 | 4.57 ± 0.65 | 6.51 ± 0.82 | 7.29 ± 0.55 | 8.06 ± 0.94 | 0.79 ± 0.08 |
| Glimepiride | 1.94 ± 0.24 | 2.01 ± 0.31 | 2.41 ± 0.57 | 2.73 ± 0.39 | 3.98 ± 0.46 | 5.66 ± 0.69 | 7.29 ± 1.06 | 0.73 ± 0.05 |
| Chloroquine | 1.68 ± 0.31 | 1.93 ± 0.51 | 2.10 ± 0.48 | 2.47 ± 0.33 | 2.98 ± 0.41 | 4.21 ± 0.70 | 5.59 ± 0.91 | 0.73 ± 0.10 |
| Carvedilol | 1.50 ± 0.15 | 1.89 ± 0.25 | 2.03 ± 0.64 | 2.61 ± 0.57 | 2.95 ± 0.58 | 3.47 ± 0.52 | 3.86 ± 1.09 | 0.72 ± 0.03 |
| Oxazepam | 0.97 ± 0.17 | 1.84 ± 0.26 | 2.43 ± 0.63 | 2.71 ± 0.63 | 3.82 ± 0.89 | 3.96 ± 0.66 | 4.17 ± 0.68 | 0.75 ± 0.06 |
| Mepacrine* | 0.50 ± 0.03 | 0.52 ± 0.00 | 0.74 ± 0.27 | 0.89 ± 0.45 | 1.49 ± 0.87 | 1.81 ± 0.94 | 2.06 ± 0.97 | 0.77 ± 0.06 |
| Proguanil* | 0.45 ± 0.03 | 0.55 ± 0.13 | 0.62 ± 0.52 | 0.87 ± 0.51 | 1.94 ± 0.46 | 2.16 ± 0.13 | 2.27 ± 1.21 | 0.74 ± 0.04 |
| Doxycycline ** | 0.39 ± 0.00 | 1.71 ± 0.26 | 1.54 ± 0.71 | 1.82 ± 0.66 | 2.18 ± 0.43 | 2.29 ± 1.0 | 2.39 ± 1.20 | 0.76 ± 0.08 |
| Mefloquine*** | 0.33 ± 0.00 | 0.35 ± 0.00 | 0.42 ± 0.13 | 0.53 ± 0.07 | 0.60 ± 0.05 | 1.07 ± 0.72 | 1.56 ± 0.93 | 0.72 ± 0.05 |
| **Control > 12.00** |  |  |  |  |  |  |  | 0.76 ± 0.05 |

A comparison of the *in vitro* activities of the four best frontrunners and the standard reference (albendazole and mebendazole) across three assay points revealed the superior activity of frontrunners. The activities of the frontrunners were significantly different from the reference standards at p<0.05 across all assay points. Asterisk sign on the bars depict the frontrunner drugs activities. This result is presented in Figs 4 and 5.

**Fig 4:**
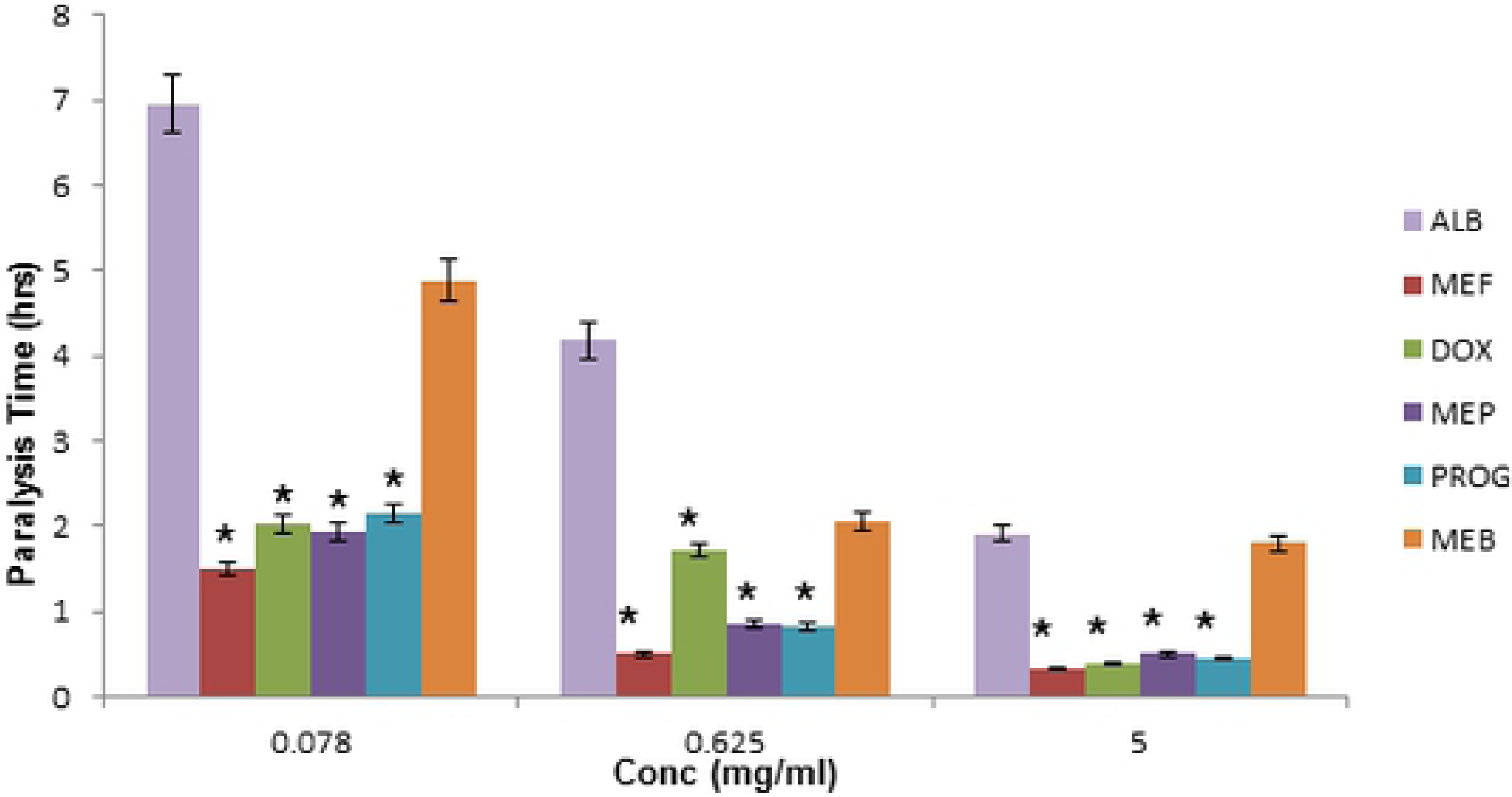
*In vitro* activity (Paralysis) of frontrunners and positive control at three assay points. *P<0.05 compared to albendazole and mebendazole.

**Fig 5:**
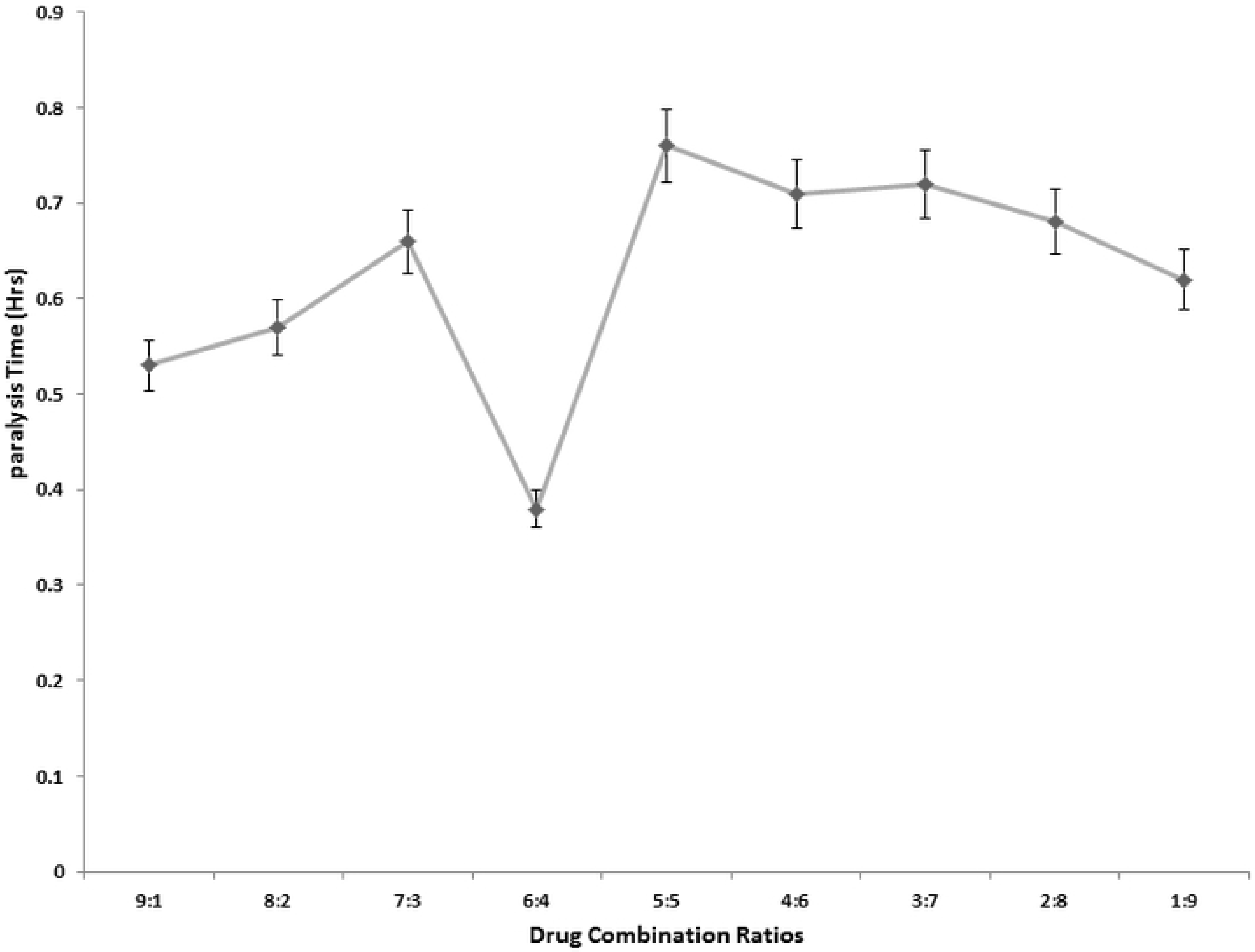
*In vitro* Activity (Mortality) of frontrunners and positive control at three assay points.. *P<0.05 compared to albendazole and mebendazole.

The synergistic testing produced the greatest activity (optimal fixed dose) at the 6:4 ratio of doxycycline and mepacrine (Fig 6 and 7). This combination produced supraadditive (synergistic) effect with paralysis combination index of 2.35 and mortality combination index of 2.23.

**Fig 6:**
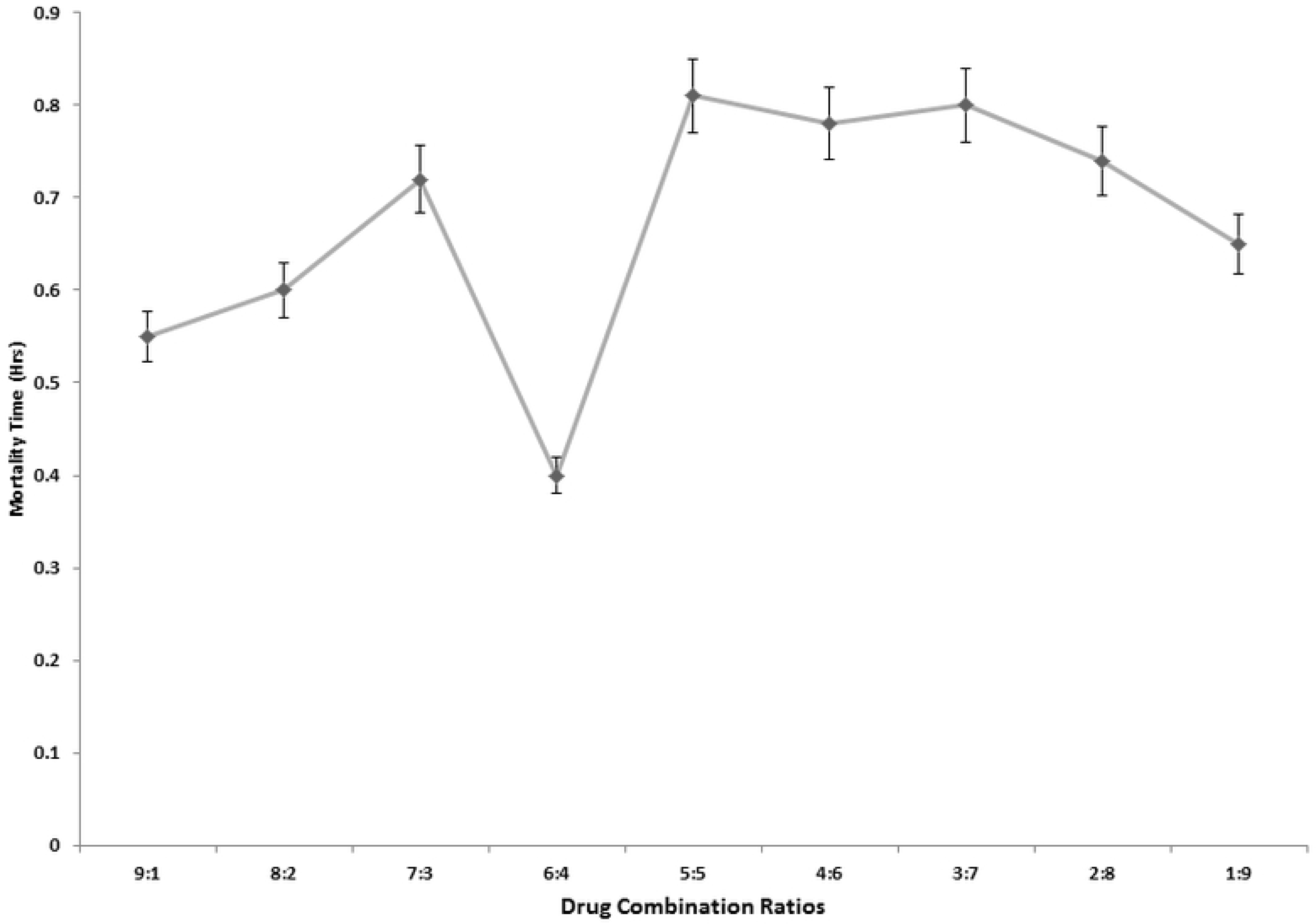
*In vitro* Paralysis Combination Studies of Doxycycline and Mepacrine.

**Fig 7:**
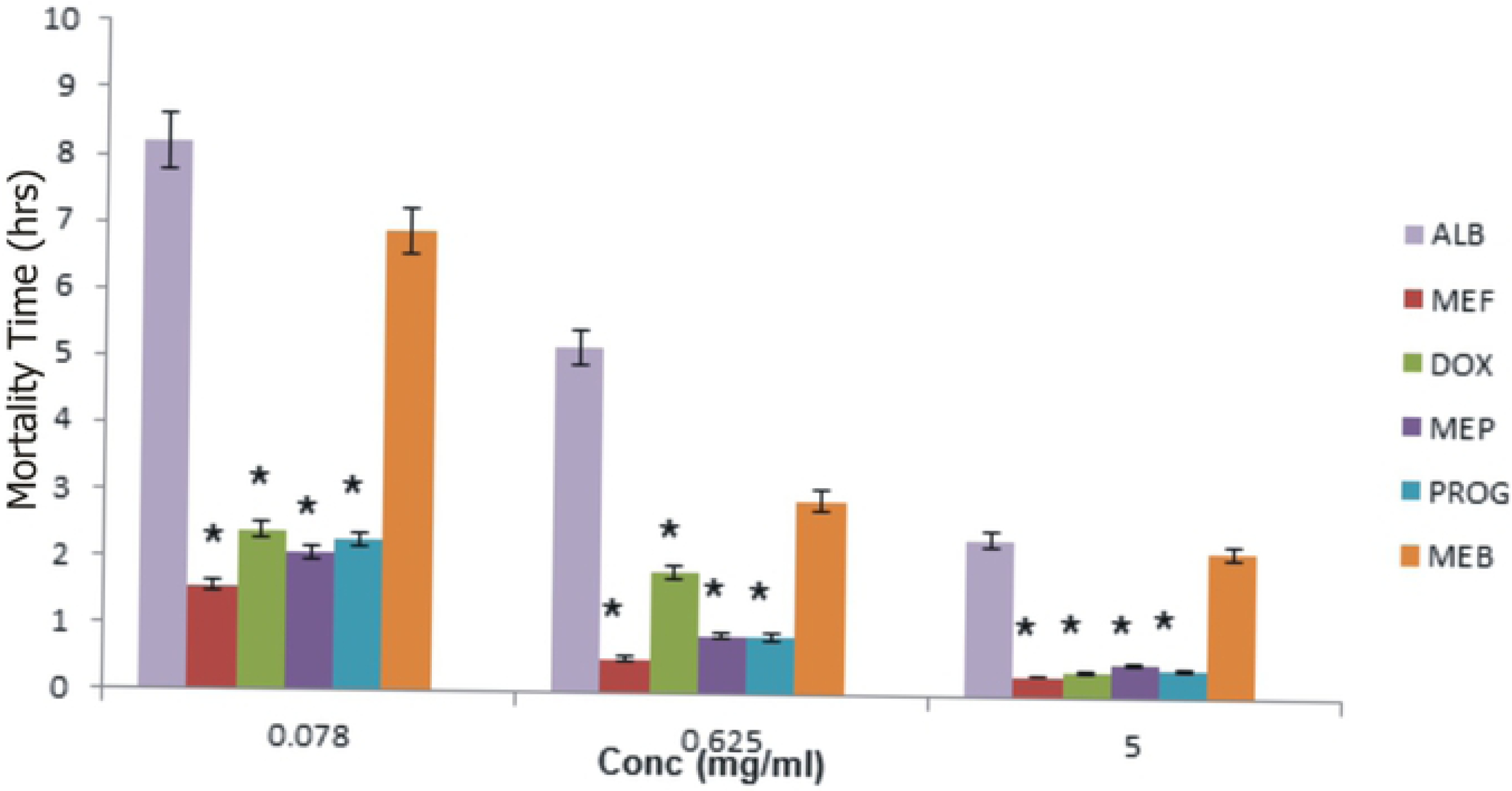
*In vitro* Mortality Combination Studies of Doxycycline and Mepacrine.

## Discussion

When drug-receptor complexes are formed, the complex with the lowest binding energy is usually considered the one with the best interaction and affinity therefore is predicted to have the best activity. The binding energies generated from docking predicted potential antihelmintic activity of some of the approved drugs that were docked.

The four major frontrunners, mefloquine, doxycycline, proguanil and mepacrine showed better binding affinities than the experimental antihelmintic compound, atpenin as well as the reference antihelmintic drugs, the benzoimidazoles.

The expectation is that these four drugs with binding energies between −11.0 to −8.1 kcal/mol will have activities that would surpass that of atpenin. Their exploitation of same binding pocket and interaction with same amino acid residues at the receptor site is suggestive of their ability to affect the receptor in the same manner as the benzoimidazoles hence, predicting the same mode of action. Doxycycline and proguanil made polar interactions with polar residues of the receptor which are expected to be of clinical significance. The critical role of hydrogen bonds in ligand binding has been stressed by Wade and Goodford [19]. The relative abundance of hydroxyl groups in doxycycline structure made for enhanced polar contacts while desolvation was pivotal to hydrophobic contacts of mefloquine and mepacrine due to disruption of the structure of bulk water leading to decreased entropy [5]. The high aqueous solubility of doxycycline is ascribed to the additional polar contacts it made with appreciable proton donors from the guanidinum group of the amino acid, arginine at the receptor binding site. These interesting *in silico* observations generated interest to test for *in vitro* activity.

The activities of the benzoimidazoles were inferior to that of the frontrunner drugs with respect to paralysis and mortality times (1.80 – 1.94 hr) as against (0.33 – 0.50 hr) thereby validating the *in silico* observations. However the high aqueous solubility of doxycycline is a drawback in application as intestinal antihelmintic chemotherapy since an ideal intestinal antihelmintic agent requires extended presence in the vicinity of the parasite to be effective. A compelling need for structural modification therefore arises which can be achieved either through introduction of halogen or aromatic groups into the structure or attachment of larger alkyl side chain to deliver insoluble derivatives that would serve as ideal intestinal antihelmintics. The combination study revealed a synergistic interaction between doxycycline and mepacrine with a combination index of 2.35 and 2.23 (paralysis and mortality) and optimal activity at a fixed dose of 6:4. This may be explained by the inhibition of the metabolism of mepacrine by doxycycline and this is of great interest given the inadequacies of single molecule therapy of antihelmintics currently in addition to the increasing wave of resistance to available treatments. The works of Keiser *et al* [8] confirmed these inadequacies and the effectiveness of combination therapy in antihelmintic chemotherapy.

In addition the merits of combination therapy even in other therapeutic landscapes have also been emphasized by several success stories reported such as the works of Knopp *et al* [10] thus this observed interaction would be of immense benefit in the development of combination therapy if *in vivo* trials support this observation.

## Conclusion

The antihelmintic potentials of mefloquine, doxycycline, proguanil and mepacrine have been demonstrated. Clinical investigations are recommended to confirm these observations and possible application in antihelmintic chemotherapy especially as fixed-dose combinations.

## Materials and Methods

i. **Selection and Preparation of Receptor** Bioinformatic mining of the Protein Data Bank (PDB) was done to identify the suitable *ascaris* MRFR 3D structure for the study. The crystal structure of the MRFR (PDB code, 3vra) was obtained from the Research Collaboration Standard Bioinformatics (RCSB) database.. The Flavin Adenine Dinucleotide (FAD) and hememolecules present in the structure and the extra subunits were deleted using Chimera v.1.9. The rest of the structure (A to D) were saved in Mol 2 and PDB file formats and further processed using Auto Dock tools v.1.5.6 in order to create PDBQT file suitable for molecular docking stimulations.
ii. **Selection and Preparation of Approved Drugs** The bioactivities of the reference compound atpenin (A5) were determined using Molinspiration online tool (www.molinspiration.com). An in-house database of approved drugs was sorted and the best two bioactivity scores of the reference compound were used to query the in-house database of the approved drugs in order to select drugs with similar bioactivity scores to the reference compound. In this instance, G-Protein Coupled Receptor (GPCR) and Enzyme Inhibitor (EI) were used for selection of approved drugs. In addition, bioactivity scores in the lower range served as negative controls.
iii. **Validation of Molecular Docking Protocol** Validation of the docking protocol was implemented by reproducing the experimental complex of the probe compound with its receptor *in silico*. The receptor in complex with the probe was obtained from the RCSB database [4] by the use of bioinformatic mining and prepared for docking simulation. The probe and all hetero-molecules were deleted with Chimera v.1.9 [14]; polar hydrogen, Kollman charges were deleted then the grid box sizes and grid space centre of 10Å were determined with MGL tools v.0.1.5.6 [11]. The probe coordinates were obtained from the 3D structure in ZINC^®^ database [17] to determine the conformation of the non-complexed atpenin prior to docking with the target. Finally, all hydrogen atoms, torsions and all rotatable bonds were allowed in their natural states. The outputs were generated in PDBQT extension. Docked conformations were visualized in PyMol v. 1.4.1 and docked poses were compared with the experimental crystal structure of the probe by superimposition of the atpenin (A5).
iv. **Molecular Docking Simulations** The molecular docking stimulations of the approved drugs and the reference compound (atpenin) were implemented using Auto Dock Vina v.4.0. The search grid was dictated by the location of the atpenin binding site on the enzyme structure.
v. **Post Docking Analysis** The post docking analysis was done to determine the bonds and the various amino acids involved in the binding between the protein and ligands. This was implemented using PyMol v.1.4.1. And the binding free energies of the best binding conformations of the complexes were obtained and recorded.

### *In vitro* analysis

*In vitro* antihelmintic evaluation was done by determination of the paralysis and mortality times of *Pheretima posthuma*(adult earthworm).

*Pheretima posthuma* was chosen as surrogate due to acute unavailability of the investigated nematodes from sacrificed livestock and its amino acid sequence similarity to those nematodes by virtue of a protein sequence alignment query.

### Worm Collection and Preparation

The earthworms were harvested from swampy soil in Agulu, Anambra State, Nigeria and were stabilized in the soil marsh from where they were scooped and kept under cold chain till time of investigation.

The worms were identified by Mrs. Olue Annastasia of the Parasitology and Entomology Department of Nnamdi Azikiwe University, Awka.

### Drugs: Purchase and Preparation

The drugs used for the bioassay were obtained from registered pharmacies in Nigeria and a few from Boots, United Kingdom.

Patent holders brands were used or brands from reputable manufacturers and seven assay points were chosen between 0.078 to 5.0 mg/ml and were prepared as stock solutions.

### Preparation of Drugs for Synergistic Testing

Doxycycline and mepacrine were selected for combination studies from among the four best frontrunners using effect-based strategy [6]. The combination of both drugs were prepared in 9 ratios of 5 mg/ml assay concentration (9:1, 8:2, 7:3, 6:4, 5:5, 4:6, 3:7, 2:8, 1:9). Paralysis and mortality effect of both drugs separately and when combined were used for the calculation of their combination index using the formula.

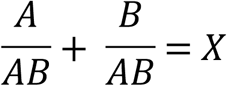

Where A = Effect of doxycycline, B = Effect mepacrine, AB = Effect of different ratio combinations of doxycycline and mepacrine. When X = 1 (additive interaction), X > 1 (synergistic interaction) and X < 1 (antagonistic interaction).

### Determination of paralysis and mortality times of frontrunners

The determination of paralysis and mortality times of frontrunners among the selected approved drugs was evaluated as described by [2].

Five worms of average weight were rinsed with distilled water and placed in 20 ml solution of each drug in a standard petri dish according to the labeled concentrations.

The petri dishes were mechanically swirled to ensure the entire worm bodies were covered in the drug solution. The worms were then monitored for paralysis and mortality times and the observations were recorded. This same procedure was replicated in the synergistic testing. Albendazole served as standard reference while distilled water was negative control. Paralysis was assessed as a situation when the worm loses muscular tone and unable to move its body except with vigorous shaking or when pricked with an object while mortality was considered when the worm does not move its body even when placed in water bath at 50 ^0^C.

### Statistical Analysis

Data collected were analyzed using the one-way ANOVA statistics with SPSS version 16.0.

